# Disease and phenotype relevant genetic variants identified from histone acetylomes in human hearts

**DOI:** 10.1101/536763

**Authors:** Wilson Lek Wen Tan, Chukwuemeka George Anene-Nzelu, Eleanor Wong, Hui San Tan, Pan Bangfen, Chang Jie Mick Lee, Matias Ilmari Autio, Michael P. Morley, Kenneth B. Margulies, Thomas P. Cappola, Marie Loh, John Chambers, Shyam Prabhakar, Roger Foo

## Abstract

Identifying genetic markers for heterogeneous complex diseases such as heart failure has been challenging, and may require prohibitively large cohort sizes in genome-wide association studies (GWAS) in order to demonstrate statistical significance^1^. On the other hand, chromatin quantitative trait loci (QTL), elucidated by direct epigenetic profiling of specific human tissues, may contribute towards prioritising variants for disease-association. Here, we captured non-coding genetic variants by performing enhancer H3K27ac ChIP-seq in 70 human control and end-stage failing hearts, mapping out a comprehensive catalogue of 47,321 putative human heart enhancers. 3,897 differential acetylation peaks (FDR < 0.05) pointed to recognizable pathways altered in heart failure (HF). To identify cardiac histone acetylation QTLs (haQTLs), we regressed out confounding factors including HF disease status, and employed the G-SCI test^2^ to call out 1,680 haQTLs (FDR < 0.1). A subset of these showed significant association to gene expression, either in *cis* (180), or through long range interactions (81), identified by Hi-C and Hi-ChIP performed on a subset of hearts. Furthermore, a concordant relationship was found between the gain or disruption of specific transcription factor (TF) binding motifs, inferred from alternative alleles at the haQTLs, associated with altered H3K27ac peak heights. Finally, colocalisation of our haQTLs with heart-related GWAS datasets allowed us to identify 62 unique loci. Disease-association for these new loci may indeed be mediated through modification of H3K27-acetylation enrichment and their corresponding gene expression differences.

## Main

Chromatin QTLs are abundant in a wide range of cell types, and contribute to phenotypic variation^3^-^9^. Genetic variants at noncoding regulatory sequences may indeed mediate their effect on complex traits and diseases through their influence on histone modifications, and downstream consequences of chromatin remodelling or gene expression^8^. Since genetic markers for more heterogeneous complex diseases such as heart failure (HF) have remained elusive, despite massive cohorts used in large GWAS analyses^1^, direct epigenetic profiling for chromatin QTLs using specific human tissue may help to prioritise variants for disease-association. Moreover, instead of the analysis for expression QTL^10^, which is easily compromised by the instability of RNA molecules in difficult-to-obtain tissue such as human hearts, the relative stability of chromatin^11^ instead offers a distinct advantage with histone QTL analysis. Hence, we performed H3K27-acetylation ChIP-seq to identify active cardiac enhancer loci, using 70 frozen post-mortem human left ventricles from Caucasians, aged 36 to 76 years old (36 end-stage HF and 34 non-failing (NF) controls) (**Supplementary Table 1**). **Fig. 1a** depicts the schematic workflow of our experimental design. We sequenced to an average of 117 ± 52 million unique paired-end reads per ChIP-seq library for each sample (**Supplementary Table 2**). Using the DFilter^12^ peak-caller, a consensus set of 47,321 H3K27ac peaks was obtained (**Extended Fig. 1**; **Supplementary Table 3**). Peak calls were analysed carefully for the quality of enrichment scores, ChIP-seq fragment reads, and inter-sample replicability (**Extended Fig. 1a, Supplementary Table 2**). As anticipated, our H3K27ac peaks also correlated well to the ENCODE dataset for human heart^13^ (**Extended Fig. 1**).

**Figure 1.**
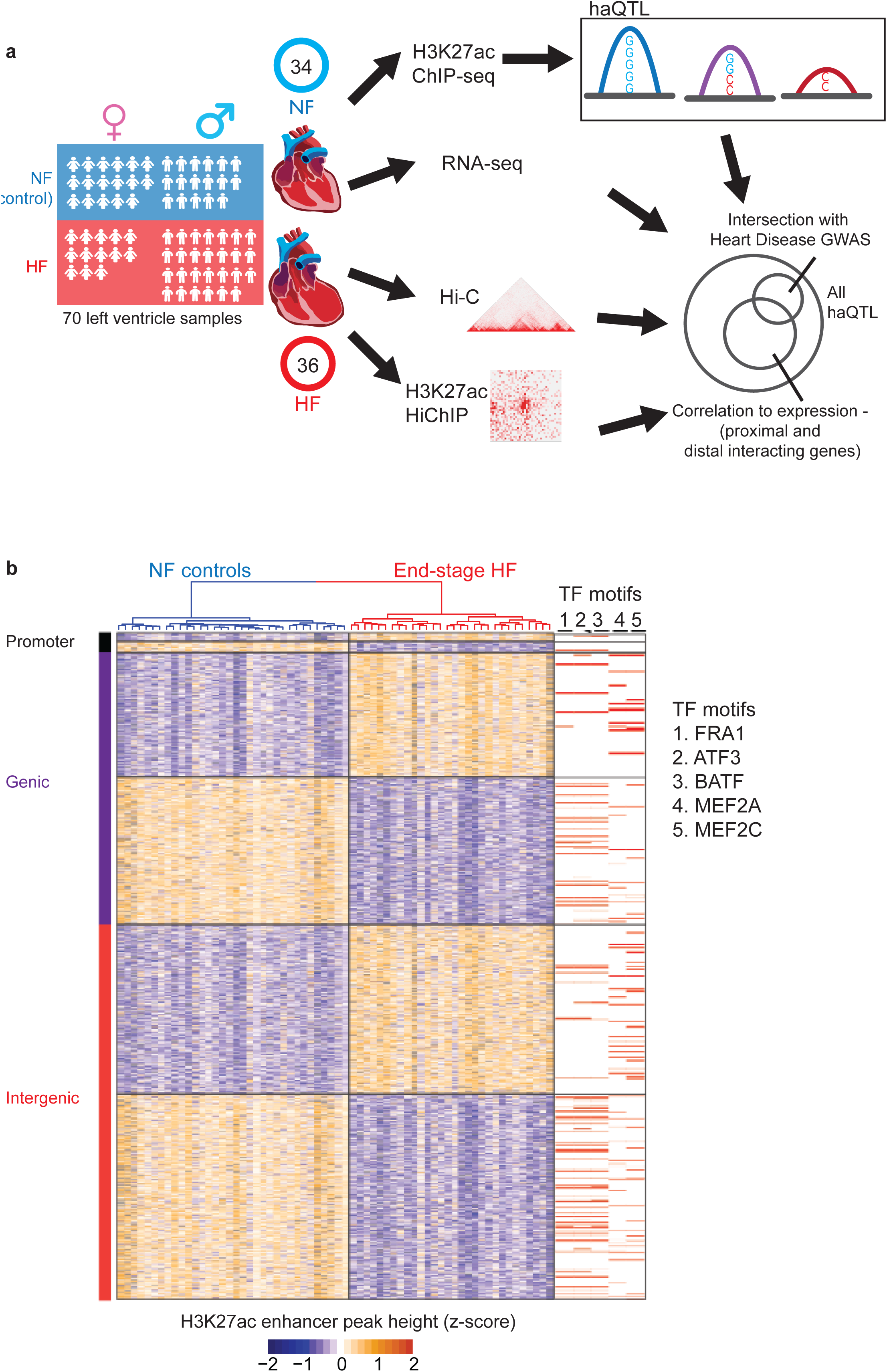
Schematic and differential H3K27-acetylation between non-failing control (NF) and failing (HF) hearts. **(a),** Schematic for the experimental workflow. ChIP-seq was performed on 70 human left ventricles (LV). G-SCI test^2^ was performed to call out haQTLs, followed by downstream analyses for correlation to gene expression and overlap with published GWAS datasets. RNA-seq was performed so as to generate matched tissue transcriptomes. Hi-C and HiChIP were performed for 2 samples to generate a cardiac chromatin connectome. **(b),** Heatmap showing differential H3K27-acetylation loci (FDR<0.05, fold-change ≥ 1.3), and transcription factor (TF) binding motifs enriched at these loci. Differential loci are categorized based on their localisation to gene promoter (black bar), intragenic (blue) or intergenic (red) regions.

To further confirm the functional validity of our acetylation peaks, we assessed for differential acetylation (DA) between control NF and diseased HF hearts. First, we normalized global peak heights for GC content (**Extended Fig. 2a**) and distributional skews. Next, we corrected for confounding effects by regressing out technical covariates, and biological covariates including age, sex, and proportion of cardiac myocytes for each sample (**Extended Fig. 2b**). Corrected peak heights were used to identify DA loci between NF and HF (Wilcoxon rank sum test; FDR < 0.05, fold change ≥ 1.3). 2,130 peaks showed significant decrease, while 1,767 peaks showed significant increase, for H3K27ac enrichment in HF (**Supplementary Table 4**). Using the full set of 3,897 DA peaks, NF and HF hearts segregated well on hierarchical clustering (**Fig. 1b**). DA loci were subjected to the Genomic Regions Enrichment of Annotations Tool (GREAT)^14^, revealing familiar heart failure pathophysiological terms of cardioblast differentiation, skeletal muscle tissue development and wnt signalling pathways in upregulated loci, and DNA damage and sarcomere organization in downregulated loci^10^,^15^ (Binomial FDR Q value < 0.05; **Supplementary Table 5**). As TFs are generally bound at regulatory loci of active enhancers as marked by H3K27ac^16^,^17^, we also analysed DA loci for the enrichment of TF motifs using HOMER (**Fig. 1b**). Key cardiac TF motifs MEF2A and MEF2C were enriched in upregulated peaks, consistent with previous evidence of their increased activity in HF^18^. AP1 and ATF3 motifs were enriched in downregulated peaks, also consistent with the reported reduced activity of ATF3 in worsening hypertensive left ventricular modelling^19^. Hence, analysis of the loci for differential H3K27ac enrichment implicated established molecular pathways of heart failure disease pathophysiology and progression.

**Figure 2.**
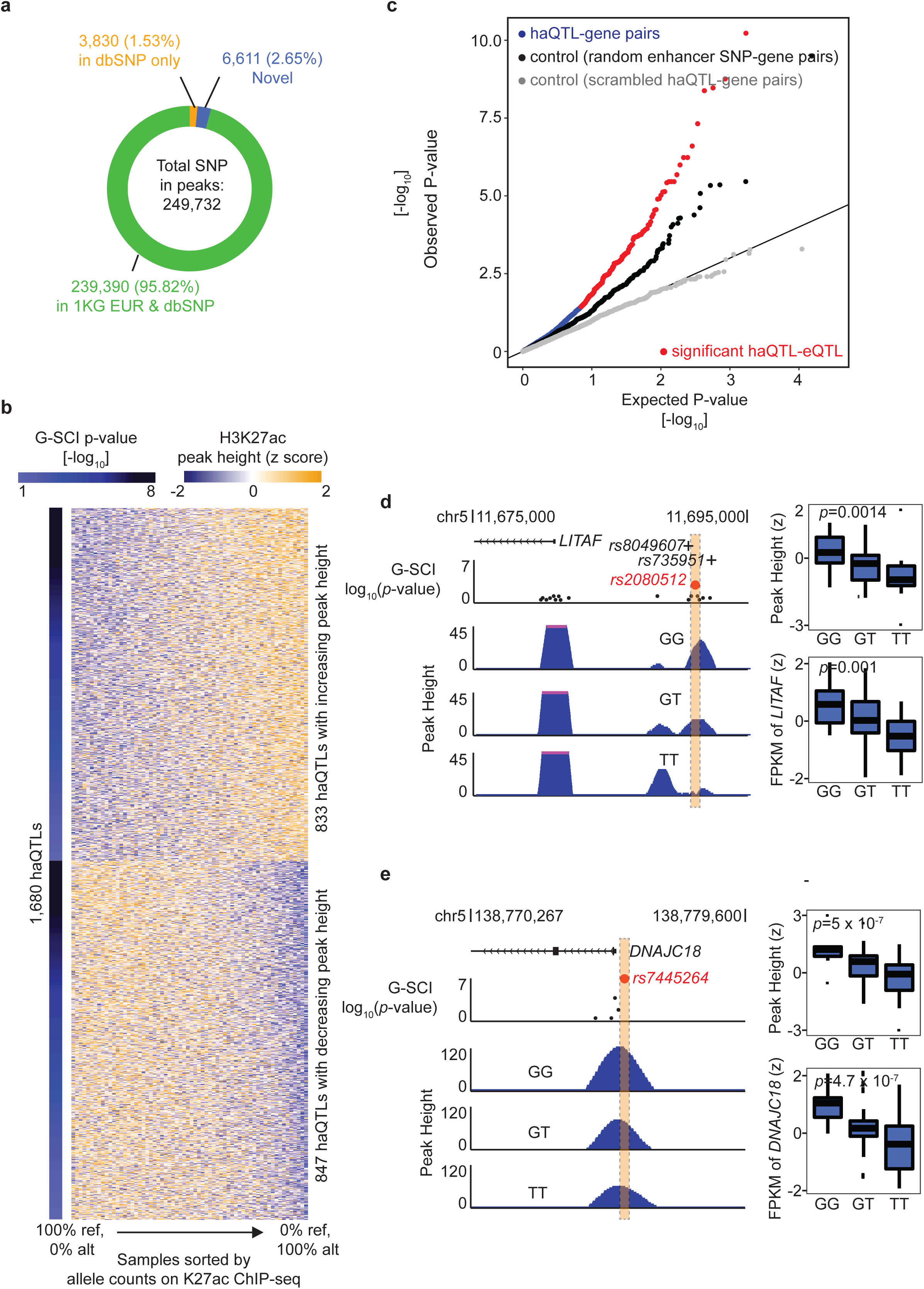
Histone-acetylation Quantitative Trait Loci (haQTL) **(a),** Pie chart showing the 249,732 SNPs underlying all H3K27-acetylation loci. **(b),** Heatmap showing 1,680 haQTLs (FDR<10%). Each row displays the H3K27ac peak heights for all LV samples at a single haQTL, arranged in order of their reference/alternative allele read count at the haQTL. The alternative allele for each sentinel SNP of 1,680 haQTLs was associated with either increasing H3K27ac peak heights (upper), or decreasing peak heights (lower). **(c),** Quantile-quantile (Q-Q) plot of the linear regression *P*-values for the association between the expression of genes (±100 kb) and haQTL, based on haQTL-gene pairs (blue), compared to random enhancer SNP-gene pairs (black), and scrambled haQTL-gene pairs where the labels between haQTL and genes were scrambled (grey). 180 *bona fide* haQTL showed gene expression associations (red) (FDR<20%). **(d),** An example of a haQTL (sentinel: rs2080512, red spot; FDR=4.29 × 10^−2^) residing in an intergenic region 13 kb upstream of *LITAF*. Black dots represent other SNPs in the enhancer locus. The alternative allele, T, is associated with reduced H3K27ac peak heights, and reduced gene expression (FDR=0.057; boxplots). This example, rs2080512, is also in linkage disequilibrium (LD) with two SNPs (rs8049607, R^2^=0.83; rs735951, R^2^=0.94, represented by +) that met genome-wide association significance for cardiac electrographic QT Interval^21^. **(e),** Another example of a haQTL (sentinel: rs7445264, red spot; FDR=5 × 10^−7^) residing in the promoter of *DNAJC18*. Samples with the alternative allele, T, show consistently reduced H3K27ac peak heights, and concordant reduced gene expression (FDR=0.0075), compared to GG homozygotes (boxplots).

Next, in order to curate for the genetic variants embedded in the cardiac H3K27ac enhancer loci, we performed variant-calling using GATK to identify SNPs within the H3K27ac peaks, arriving at a final high quality set of 249,732 SNPs (**Fig. 2a**). Nearly all SNPs (97.35%) were as annotated before in 1K Genome or dbSNP, whereas 6,611 novel SNPs were sufficiently well covered by sequencing read counts to give confidence of their validity (**Extended Fig. 3a**). Moreover, we also validated a handful of SNPs using the orthogonal method of Sanger sequencing (**Extended Fig. 3b, c**). We then proceeded to use the G-SCI test^2^, to analyse for SNPs that correlated to their local *cis* acetylation peak heights (i.e. histone acetylation QTLs, or haQTLs). Importantly, in order to make sure that the general disease effect is removed from the effect of regulatory SNPs, we adjusted acetylation peak heights by regressing out the HF disease status (**Extended Fig. 2c**). 1,680 H3K27ac peaks were finally identified to have significant association with at least one SNP at the same locus (10% FDR) (**Fig. 2b**, **Supplementary Table 6**). At each of these 1,680 acetylation peaks, a sentinel SNP was nominated to represent the haQTL, based on FDR ranking. The MAF of these are represented in **Extended Fig. 3d**.

**Figure 3.**
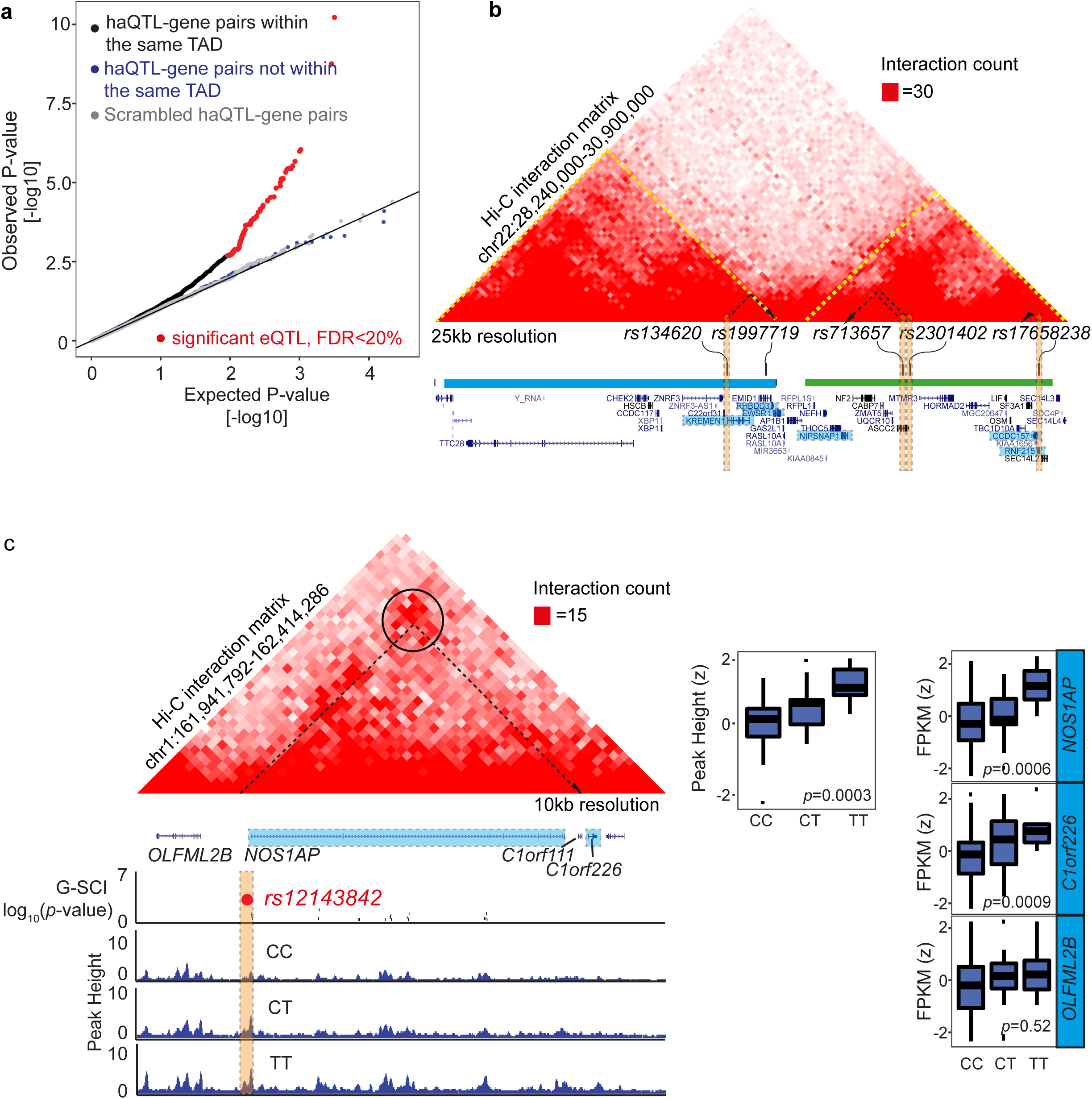
Distal gene expression associated to haQTL are confined to within topologically associated domains (TADs) **(a),** QQ plot of linear regression *P*-values for the association between gene expression and haQTLs, based on haQTL and distal gene pairs residing within the same TADs (black), compared to pairs spanning across neighbouring TADs (blue), and scrambled haQTL-gene pairs (as randomised in Fig 2c). 57 haQTLs with gene expression associations lying within the same TAD (red) achieved FDR<20%. **(b),** Snapshot showing 4 haQTLs (orange vertical boxes), one residing in an upstream TAD (blue horizontal bar), and 3 residing in the downstream TAD (green bar). TADs are also delineated by yellow dotted lines in the Hi-C interaction matrix. Each haQTL is correlated to the expression of proximal and distal genes localised to only within the same TAD. For example, rs134620 is associated with the expression of *KREMEN1*, and distal interacting genes *RHBDD3* and *EWSR1*, located 200kb away but within the same TAD. rs713657 and rs2301402 are both correlated to the expression of a distal gene *NIPSNAP1* 235 kb away, again within the same TAD. Correlated genes are coloured in blue, distal interactions are represented by dotted black arrows. **(c),** An example of a haQTL with correlated distal gene expression. rs12143842 (red spot) located in the *NOS1AP* gene promoter, is correlated to its expression (FDR=0.14), and also to the expression of a distal interacting gene, *C1orf226*, located 500kb downstream (FDR=0.10). A chromatin loop (black circle) in the Hi-C interaction matrix highlights the interaction (dotted black arrow) connecting the 2 distal loci. See Supplementary Fig. 9d for replication of this distal interaction by HiChIP. Importantly, there was no expression correlation for this SNP to its upstream gene (*OLFML2B*) that was lacking Hi-C evidence for interaction, despite being in closer proximity. rs12143842 is also a significant GWAS hit for QT interval^71^.

As haQTLs, localising to either gene promoters or intergenic distal enhancers, and correlating to enhancer peak heights, may also correlate to proximal gene expression ^8^, we overlapped the list of haQTLs with human left ventricle eQTL mapping data from the GTEx project^20^. We found 167 haQTLs in strong linkage disequilibrium (R^2^>0.8) with GTEx eQTL (**Supplementary Table 7**). By random permutation analysis, haQTLs were more enriched for GTEx eQTL, compared to non-haQTL enhancer SNPs (20-fold enrichment, empirical *P*-value < 0.001; **Extended Fig. 4**). Concurrently, we also generated RNA-seq data from our full set of left ventricles. A substantial number (180, 10.8%) of haQTL also bore expression correlation to their proximal genes (±100 kb), using our RNA-seq data at the more inclusive FDR threshold of 20%, with 40% overlap to the set from GTEx eQTL (**Supplementary Table 7**). A QQ plot (**Fig. 2c**) showed haQTL to be more enriched for expression correlation at lower association *P*-value, compared to random enhancer SNPs, or to scrambled haQTL-gene pairs where haQTL labels were randomly swapped, strengthening the notion as before that haQTL significantly associates with proximal gene expression more than by random chance. For example, **Fig. 2d** shows a haQTL located at the promoter region of *LITAF* (sentinel: rs2080512). The presence of alternative alleles correlated to reduced enhancer peak heights, and a corresponding decrease in *LITAF* gene expression. *LITAF* has been associated with cardiac electrocardiographic QT elongation with variants at the locus (LD R^2^>0.8) that met genome-wide significance in a previous GWAS^21^ (**Fig. 2d**). LITAF regulates cardiac excitation by acting as a molecular adaptor, linking NEDD4-1 ubiquitin ligases to the cardiac L-type calcium channel, and *LITAF* knockdown prolonged action potential duration and increased cytosolic calcium^22^. Similarly, for another haQTL located in the *DNAJC18* promoter (sentinel: rs9573330; **Fig. 2e**), alternative alleles also correlated with decreased peak heights, and reduced *DNAJC18* gene expression. The DNAJ/HSP40 family of proteins consists of molecular chaperones involved in cellular functions of protein folding, translation and translocation^23^. While there is little information about *DNAJC18* in the heart, some members of the family have shown importance in the regulation of cardiac potassium ion channels^24^ and mitochondrial biogenesis^25^.

**Figure 4.**
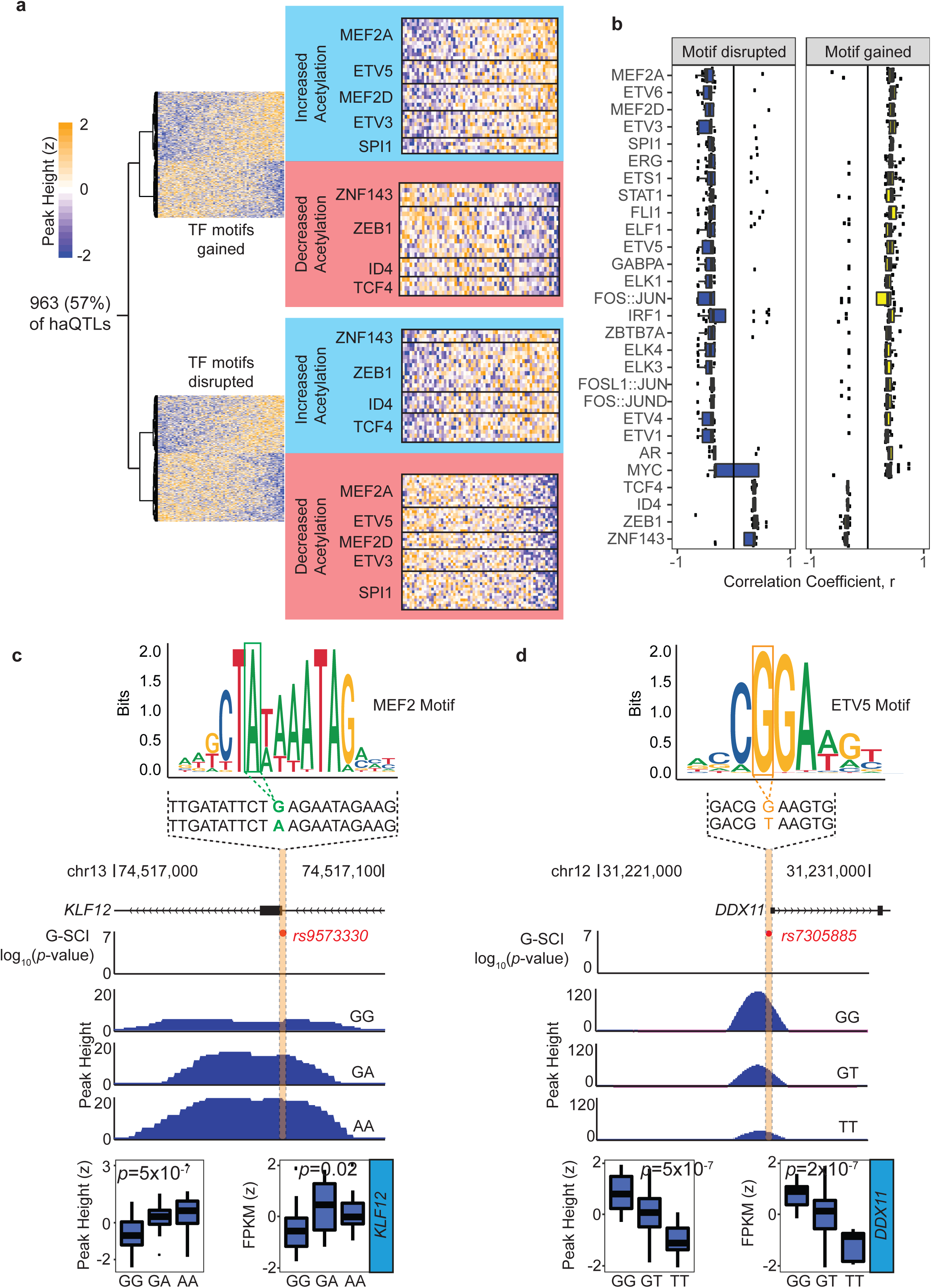
haQTLs inferred to alter transcription factor (TF) binding. **(a),** Alternative variants for sentinel SNPs at 963 (out of 1,680) haQTLs are inferred to alter at least 1 TF binding motif for cardiac expressed TFs. Heatmaps show the correlation between TF motif gained or disrupted, to the change in H3K27ac peak height. Examples of 2 unique sets of TFs (in blue and red boxes) show a converse relationship where motif gain either correlated with increased or decreased acetylation peak height, and vice versa. **(b),** Boxplots showing the association between TF motif disrupted or gained with the direction of H3K27ac peak height change. This unique set of 28 TFs show significant opposite effect between TF motif alteration and peak height change (Wilcoxon Rank Sum Test, FDR<10%). 24 TFs show increased peak height (haQTL correlation coefficient, r>0) with *de novo* motif gain, and reduced peak height with motif disruption (r<0). 4 TFs show the opposite effect of increased peak height with motif disruption, and decreased peak height with motif gain. **(c),** An example of haQTL (sentinel: rs9573330) inferred to produce a gain in MEF2A motif at a H3K27ac locus in the intron of *KLF12* (FDR=1.26 × 10^−4^). The motif gain is associated with increased H3K27ac peak height, and concordant increase in *KLF12* gene expression (FDR<20%). **(d),** An example of haQTL (sentinel: rs7305885) inferred to disrupt ETV5 motif in the promoter of *DDX11*. The alternative allele, T, is associated with reduced acetylation (FDR=1.26 × 10^−4^), and reduced *DDX11* gene expression (FDR=1.54 × 10^−4^).

Enhancers may also regulate distal gene expression over long genomic distances by long-range interactions resulting from chromatin folding^7^,^26^. We therefore also examined the association between haQTLs and the expression of their putative distal interacting genes, after building up a cardiac chromatin interactome using *in situ* Hi-C that we performed in 2 hearts (**Supplementary Table 2**). Hi-C libraries were sequenced to 540 million unique reads per sample, producing a chromatin contact matrix at 40 kb resolution. Topologically associated domains (TADs) are defined as self-interacting chromatin domains whose boundaries are enriched for insulator proteins such as CTCF and Cohesin, and whose composition have been shown to confine gene regulation via enhancer-promoter interactions to within TADs^26^,^27^. As previously suspected ^27^,^28^, our 2 Hi-C libraries showed global similarity for TADs between hearts (**Extended Fig. 5a; Supplementary Table 8**). Moreover, in a separate study, we have also found that, despite hallmark and characteristic gene expression changes in a mouse model of heart failure, the cardiac CTCF-based TAD chromatin architecture remains robust between healthy and failing hearts, in marked contrast to the alternative model of cardiac-specific CTCF knockout (Lee et al, *in press*). We also observed that our TAD boundaries were marked by characteristic epigenetic marks including CTCF^27^ (**Extended Fig. 5b-d**) and were highly insulated^29^ (**Extended Fig. 5e, f**). As anticipated therefore, we observed that a correlation between haQTL and distal gene expression were more likely to occur for haQTL-gene pairs that resided in the same TAD, as opposed to haQTL-gene pairs spanning across TADs, or scrambled haQTL-gene pairs (**Fig. 3a, b**). 65 associations (involving 57 haQTLs) for distal haQTL-gene pairs within the same TAD (at least 10 kb apart) met the FDR threshold of 20% (**Supplementary Table 7**). For example, **Fig. 3c** shows the associations for sentinel SNP rs12143842, in the promoter locus of *NOS1AP*. The alternative allele associated with increased enhancer peak height, and increased expression of the proximal *NOS1AP* gene and a distal interacting gene *C1orf226*, but not the closer *OLFML2B* gene which was in a neighbouring TAD. While the function of *C1orf226* is unknown, NOS1AP is a known regulator of neuronal nitric oxide synthase (NOS1), which is highly expressed in the mammalian heart and regulates cardiac excitation-contraction coupling and repolarization ^30^.

**Figure 5.**
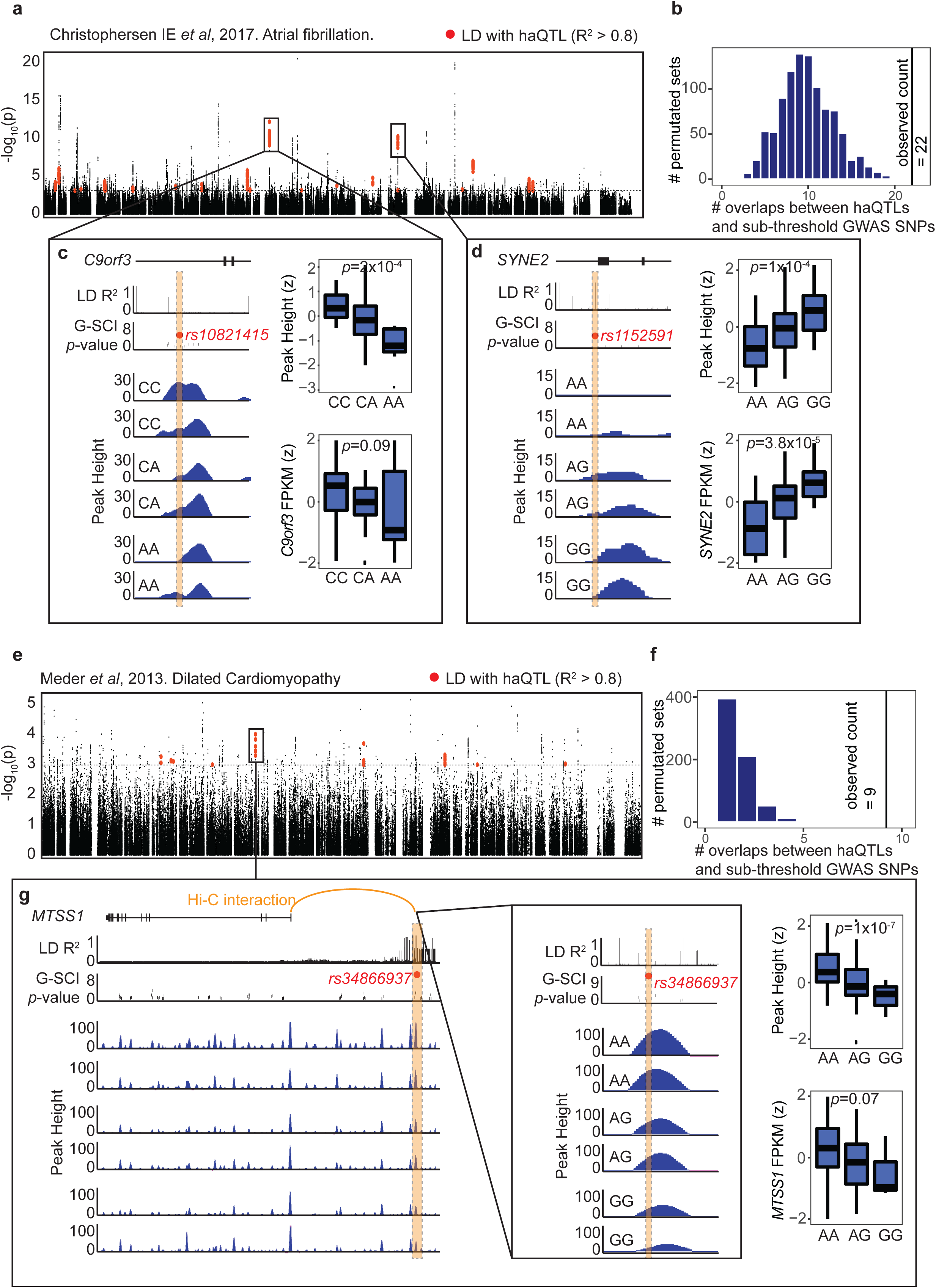
haQTLs enriched for GWAS sub-threshold SNPs. **(a),** A Manhattan plot of adjusted *P*-values for all SNPs from the GWAS meta-analysis for atrial fibrillation^45^, in LD (R^2^>0.8) with SNPs in our 249,732 cardiac enhancers (black). Red represents colocalised SNPs in 22 loci that are either in LD or overlapping with haQTLs. **(b),** 1,000 permutation sets of 1,000 enhancer SNPs, matching for distribution to nearest TSS distance, and MAF, to simulate the expected number of overlap between GWAS and haQTLs. We observed 22 colocalisation between haQTL and atrial fibrillation GWAS SNPs, a 2-fold enrichment from the expected (empirical *P*-value<0.001). **(c, d),** Examples of 2 independent haQTLs (sentinels: rs10821415 and rs1152591) colocalised with atrial fibrillation GWAS sub-threshold SNPs. For (**c)**, the alternative A allele for rs10821415 is associated with reduced H3K27ac peak height (FDR=0.0105) located within a *C9orf3* intron, and a trend of reduced gene expression FDR=0.74) (boxplots). Notably, rs10821415 is a significant eQTL for *C9orf3* in the GTEx project ^20^. For (**d)**, the alternative G allele for rs1152591 is associated with increased H3K27ac peak height (FDR=0.0006) located within a *SYNE2* intron, and increased *SYNE2* gene expression (FDR=0.018) (boxplots). **(e),** A Manhattan plot of adjusted *P*-values for all SNPs from the GWAS for dilated cardiomyopathy^49^, in LD (R^2^>0.8) with our enhancer SNPs (black). Red represents colocalised SNPs in 9 loci that are either in LD or overlapping with haQTLs. **(f),** 1,000 permutation sets of 1,000 enhancer SNPs, matching for distribution to nearest TSS distance, and MAF, to simulate the expected number of overlap between GWAS and haQTLs. We observed 9 colocalisation between haQTL and dilated cardiomyopathy GWAS SNPs, a 9-fold enrichment from the expected (empirical *P*-value<0.001). **(g),** An example of haQTL (sentinel: rs34866937) colocalised with dilated cardiomyopathy GWAS sub-threshold SNPs. The alternative G allele is associated with reduced H3K27ac peak height (FDR=1.26 × 10^−4^) located 70 kb from the promoter of *MTSS1*, and a trend of reduced *MTSS1* gene expression (FDR=0.71) (boxplots). Again, rs34866937, is a significant eQTL for *MTSS1* in the GTEx project ^20^.

Since it would be of value to assign haQTL to more distal interacting genes, we also deepened our study further by constructing and sequencing H3K27ac HiChIP libraries from 2 heart samples, enabling us to identify chromatin interactions based on H3K27ac enhancers at the anchors^31^. On average, 64 million unique valid read-pairs were sequenced per HiChIP library. Since both HiChIP libraries were again highly correlated (**Extended Fig. 6a)**, we combined both replicates to attain the greater depth of 194 million unique di-tags (388 million reads). Using HICCUPS, 14,698 H3K27ac based chromatin loops were called (10% FDR; **Supplementary Table 9; Extended Fig. 6**). H3K27ac enrichment at loop anchors was confirmed, with 82% overlapping to H3K27ac ChIP-seq peaks, in at least 1 of the anchors (**Extended Fig. 6c-e**). 613 of these loops were anchored by a haQTL. The majority of haQTL-containing anchors (407/613, 60%) interacted with other H3K27ac-enriched loci ∼30 kb away, and 45% (184/407) also contained expressed gene TSS. From these, we recognised 15 haQTLs (FDR 20%) that we had already noted to bear distal gene expression association, as identified in **Fig. 3a**. The analysis making use of the HiChIP connectome yielded a further 24 unique loci, where the distances between the haQTL and genes were greater than 100 kb apart (**Supplementary Table 7**). Specific examples for these haQTLs with distal gene expression association are shown in **Extended Fig. 7**.

Binding of TF to genomic regulatory regions such as enhancers is important for transcriptional regulation^17^, and specificity of TF binding often depends on the recognition of unique DNA motif sequences^17^. Functional genetic variants located at enhancers may therefore be inferred to disrupt or create TF DNA motifs, thereby changing TF binding affinity, and in turn influencing histone acetylation, and corresponding gene expression differences^5^,^8^,^32^. Using motifbreakR^33^, we therefore analysed our haQTLs for putative disruption or gain of TF motifs, based on the reference and alternative alleles at each locus (**Supplementary Table 10**). Keeping the relevance only to TF that are cardiac expressed based on our RNA-seq dataset, we inferred that TF binding motifs were altered by alternative alleles (**Fig. 4a**). By comparing to 1,000 permutated SNP sets of non-haQTL, we validated that haQTLs were significantly enriched for TF motif alteration (**Extended Fig. 8a, b**). Altogether, 963 loci were inferred to have either a *de novo* gain in TF motif, a loss in TF motif, or had SNP variants at the same locus that produced gains and losses for 2 or more different TF motifs. In analysing for the correlation between motif gain or disruption and acetylation peak heights, we further noticed a specific set of TFs that showed a remarkable converse relationship (**Fig. 4a, b**). Gain of the MEF2 family motif (MEF2A and MEF2D), as well as that for ETV5, ETV3, SPI1 and others, was associated with increased acetylation peaks, whereas disruption of the same TF motifs was associated with reduced acetylation. As examples, **Fig. 4c, d** show the gain and loss of MEF2 and ETV5 motifs at the *KLF12* and *DDX11* loci, respectively, associated with the increase or decrease of enhancer peak heights, and corresponding gene expression changes. The KLF family of transcription factors are DNA-binding TFs that regulate fundamental cellular processes such as growth, proliferation and apoptosis, and KLF family members are known to regulate cardiac hypertrophy in response to external stimuli ^34^. Conversely, a gain in motif for the second set of TFs: ZNF143, ZEB1, ID4 and TCF4, was associated with decreased acetylation peaks, and vice versa. This is interesting because it implies that in collaboration with the H3K27-acetylation and de-acetylation machinery, inferred TF binding differences at these loci appear to be sufficient for concordant correlations to K27-acetylation peak heights, suggesting a primary functional role for the TFs at haQTLs. The 2 sets of TFs may promote or repress the active enhancer H3K27ac mark, respectively. As also found in the foregoing analysis of DA loci, MEF2 TFs are important signal-responsive mediators in the transcriptional reprogramming of cardiac development and disease^15^,^35^. They are known to bind to and recruit the histone acetyl-transferase p300 in cardiac cells ^36^. Previous studies show that MEF2 overexpression leads to dilated cardiomyopathy as MEF2 binds to the promoters of target genes to activate a fetal gene programme^37^. Similarly, ETS TFs (ETV3, 5, 6, SPL1, ERG, ETS1, FL1) are important in cardiac development ^38^, and also recruit the histone acetyl-transferase p300^39^. Conversely, TCF4 appears to be a negative regulator of gene expression ^40^, involving an interaction with histone deacetylases (HDACs)^41^. ID4^42^ and ZEB1^43^ are also known repressors of gene expression. Like TCF4, ZEB1 recruits HDACs at target gene promoters^44^. Other examples of motif changes at haQTLs are shown in **Extended Fig. 8c, d**. Globally, we observed that motif changes at haQTL with altered H3K27-acetylation also correlated concordantly to gene expression changes (**Extended Fig. 8e**). All together, these findings suggest that altered TF binding affinity inferred from variants in haQTL may lead directly to enhancer histone acetylation differences and changes in the expression of cardiac phenotype-driver genes.

Finally, in order to apply haQTLs to prioritizing variants for disease-association, we sought to colocalize our 1,680 haQTL against published heart-related GWAS. For these, we chose the large meta-analysis performed for atrial fibrillation^45^, studies from the EchoGen consortium on cardiac structure and function^46^, CHARGE studies performed for incident heart failure^47^, and heart failure mortality^48^, and a three-staged GWA study performed on 4,100 dilated cardiomyopathy cases and 7,600 controls^49^. Using an inclusive *P*-value cutoff of 0.001 for sub-threshold GWAS SNPs^50^, we narrowed down further to those that lie in cardiac active enhancer loci as marked by H3K27-acetylation, and intersected them with our catalogue of haQTLs or SNPs in LD with our haQTLs (**Fig. 5**; **Extended Fig. 9**). For each GWAS dataset, we performed independent permutation analyses using random enhancer-localised SNPs that matched our haQTLs for MAF and distance to gene promoters. Indeed, a 2 to 9 fold enrichment was observed for colocalized variants across the 10 GWAS datasets, confirming that our haQTLs were significantly enriched for heart disease-associated variants, compared to 1,000 sets of 1,000 randomly permutated enhancer-localised SNPs. Twenty-two loci were found in overlap between haQTLs and hits from the atrial fibrillation GWAS ^45^. Two of the top loci (sentinels: rs10821415 and rs1152591; **Fig. 5a-d**) showed correlation to acetylation peak heights at intragenic enhancers, and also correlation to *cis* gene expression of *C9orf3* and *SYNE2*, respectively. *SYNE2* (Nesprin-2) and other nesprin isoforms, are spectric-repeat proteins, highly expressed in cardiac muscle, important for maintaining the integrity of the nuclear envelope by coupling the nucleoskeleton to the cytoskeleton ^51^. *SYNE2* mutations cause muscular dystrophy and dilated cardiomyopathy ^51^. The function of *C9orf3* is as yet unknown. **Fig. 5e-g** shows another example where the sentinel SNP rs34866937, residing in an intergenic locus, was one that overlapped between haQTL and the GWAS for dilated cardiomyopathy^49^, as well as the EchoGen GWAS for cardiac structure and function^46^. Our Hi-C matrix revealed a chromatin loop connecting the intergenic enhancer locus to the *MTSS1* gene promoter ∼120 kb away, which we also confirmed by 3C-PCR. The alternative allele at this locus was correlated to reduced H3K27ac peak height, and reduced expression of *MTSS1* (**Fig. 5g**). Remarkably, an *Mtss1*-knockout mouse has been generated based on the prioritisation of another variant (rs12541595) in this distal interacting enhancer locus (Morley et al, *in revision*), also because of its association from in the EchoGen study^46^. The *Mtss1*-knockout mouse indeed shows improved cardiac function, while luciferase and CRISPR edit assays proved that it was the our sentinel SNP, rs34866937, and not the other SNPs in the cluster at the enhancer locus, that associated with *MTSS1* expression. **Supplementary Table 11** lists the 62 loci which were identified as haQTL with intersection for the same locus in at least one of the GWAS datasets.

In summary, our analysis has identified a set of 1,680 enhancer loci whose H3K27-acetylation enrichment was associated to unique genetic variants underlying each peak. A substantial number of them also bore correlation to gene expression abundance, sometimes to distally interacting genes that are hundreds of kilobases away, but nonetheless identifiable from cardiac Hi-C and HiChIP based chromatin connectome maps. Population genetic variants at noncoding enhancer loci may indeed have functional value for disease and phenotype if they perturb TF binding motifs, leading to chromatin and gene expression differences^8^. Hence, it may be speculated that some such differences are silent in the relevant tissue, until the onset of pathogenic stimuli when because of the noncoding genetic variant, a differential disease response ensues. This mechanism likely underpins an important inter-relationship between genetics and epigenetics for their relevance to disease and phenotypes. The haQTL dataset here should now prove useful for prioritising more genetic variants from other heart-related GWAS. The approach of chromatin QTL and 3D connectome analyses in disease-relevant tissue promises not just to resolve the identity of functional genetic variants, but target genes with correlated expression changes may be implicated to represent important pathways for new disease therapy.

## Acknowledgement

This work was funded by the Biomedical Research Council, Agency for Science, Technology and Research (ASTaR) Special Positioning Fund (SPF2014/004), and Individual Research Grants and Clinician Scientist Award from National Medical Research Council of Singapore. We thank Arima Genomics for use of their Hi-C kits. We also thank Dr Wenjie Sun for her advice with haQTL methodology.

