## Supplementary Methods and Figures for "Disease and phenotype relevant genetic variants identified from histone acetylomes in human hearts"

### MATERIALS AND METHODS

#### Ethical statement and human heart tissue.

The Myocardial Applied Genomics Network (MAGNet; www.med.upenn.edu/magnet), collects and banks human cardiac tissue for genomic research. All subjects or next of kin provided written informed consent for tissue donation, and analyses and all study protocols were approved by relevant institutional review boards. Left ventricular (LV) free-wall tissue was harvested at the time of cardiac surgery from subjects with heart failure undergoing transplantation and from unused donor hearts with apparently normal function. Hearts was perfused with cold cardioplegia prior to cardiectomy to arrest contraction and prevent ischemic damage, and tissue specimens were frozen in liquid nitrogen. LV sample metadata are provided in Supplementary Table 1.

#### Human cardiac H3K27ac ChIP-seq

For each ChIP-seq library, 70 mg of snap frozen human heart left ventricle tissue was cut and pounded into fine powder in liquid nitrogen. Tissues were washed twice in cold PBS (Takara, 1x protease inhibitor), cross-linked with formaldehyde (1%) for 5 min at room temperature and subsequently quenched with glycine (125mM) for 5 min at room temperature. Cell pellets were rinsed twice with cold PBS. Cells were lysed in lysis buffer (50mM HEPES-KOH, pH 7.5, 150mM NaCl, 1mM EDTA, 1% Triton X, 0.1% Sodium deoxycholate, 0.1% SDS, Takara 1X protease inhibitors) using a glass douncer tight pestle 'A' with 10 to 15 strokes to release nuclei. Centrifuge was performed at 4000 rpm for 10 min at 4ºC to collect the nuclei pellet. Nuclei were checked under microscopy for each preparation. Nuclei were lysed in nuclei lysis buffer (50mM HEPES-KOH, pH 7.5, 150mM NaCl, 1mM EDTA, 1% Triton X, 0.1% Sodium deoxycholate, 1% SDS, Takara, 1x protease inhibitor) and sonicated with Bioruptor sonicator to obtained chromatin fragments between 200 to 500 bp. Sheared chromatin was immunoprecipitated with 5 µg of H3K27ac antibody (Abcam, ab4729) with 50 µl of protein G beads (Invitrogen) overnight at 4ºC. Beads were washed and eluted in 200 µl of elution buffer (50mM Tris-HCl, pH7.5 and 10mM EDTA), de-crosslink at 65ºC overnight. Pulldown was phenol/chloroform treated and DNA purified by ethanol precipitation. Library preparation was performed using 2 ng ChIP DNA with NEB ultra II library preparation kit, according to manufacturer's protocol. 10-12 PCR cycles were performed using indexed primers and libraries size between 300 to 500 bp were selected. The libraries underwent paired end sequencing at 2 x 100 bp read length on Illumina Hiseq 2500 platform. Samples were processed in batches as enumerated in Supplementary Table 1. We also downloaded H3K27ac ChIP-seq data (Experiment Accession: ENCSR557DFM) from ENCODE[1].

**Read Alignment and Peak Calling**

Paired-end sequencing reads were aligned to human genome (*hg19*) genome using BWA mem version 0.7.5[2]. Duplicated reads were filtered out using Picard MarkDuplicates function. The mapped reads were fed into peak-caller, Dfilter[3], to identify significant peaks using the following setting: -ks=60, -bs=100, -lpvalue=8. Each of the ChIP-seq replicates were assessed for their Duplication Rate, Fraction of Reads in Peaks (FRiP), average correlation with other replicates and SSD. Peaks detected in at least 4 individuals were merged across samples (overlap>4) to define a consensus set of 47,321 peaks. All subsequent analyses were performed based on this 47,321 consensus peak set.

**Peak Height Normalization**

Human genome was divided into 100 bp bin. For each libraries, reads were counted in 100 bp bins, and scaled to normalize for sequencing depth. Binned counts were then adjusted by normalizing their GC-content against the average GC-content of all libraries. In each peak region, the sum of bin-wise normalized counts was defined as the peak height. In order to reduce technical variation, the peak heights in the consensus peak set were quantile-normalized [4].

### Removal of Confounding Factors

Firstly, the normalized peak heights were $log_2$-transformed. Principal-component analysis (PCA) was performed to identify potential confounding factors by correlating the top 5 principal components (PCs) with the biological covariates (etiology, conditions such as atrial fibrillation, hypertension, diabetes and ventricular tachycardia/ventricular fibrillation, age, gender, height, weight, cardiomyocyte purity) and technical covariates (sequencing batches, median library fragment insert size from paired-end reads, proportion of duplicated reads, sequencing depth and number of peaks per ChIP-seq library) (Extended Fig. 2b, c). Cardiomyocyte purity was estimated using FAC-sorting (see below). For differential acetylation (DA) analysis between end-stage heart failure and non-failing hearts, covariates that were regressed out were age, gender, height, weight, cardiomyocyte purity, and technical covariates (Extended Fig. 2b). For identifying haQTL, we regressed out all technical and biological covariates, crucially including etiology, conditions such as atrial fibrillation, hypertension, diabetes and ventricular tachycardia/ventricular fibrillation (Extended Fig. 2c). PCA was performed again after regression to confirm that no confounding factors correlated strongly with the top 5 PCs. G-SCI test analysis to identify haQTL were based on the peak height matrix after covariate regression.

### Differentially Acetylation (DA) analysis between Heart Failure (HF) and control non-failing hearts (NF)

Using the normalized peak height matrix with confounding factors removed, DA peaks were identified with the following criteria (fold-change > 1.3; FDR < 0.05; Wilcoxon rank sum test;

Benjamini-Hochberg correction). Differentially hyper-acetylated and hypo-acetylated, in bed format, were supplied to the GREAT analysis tool for pathway analysis separately using the default parameters (FDR < 0.05) [5]. For motif enrichment analysis, we used the HOMER findMotifsGenome.pl script [6]. Motif models were drawn from the TRANSFAC vertebrate database [7] and the analysis was performed separately on increased and decreased DA peaks.

**SNP-Calling Pipeline**

ChIP-seq reads were passed to the multi-sample SNP-calling pipeline. Reads used for SNP calling were de-duplicated and retained only if they were mapped to the genome in the proper orientation. We performed indel realignment, base-quality-score recalibration and SNP calling using GATK version 2.7-2 [8]. 576,482 SNPs within peaks were called using GATK Unifiedgenotyper at a SNP quality threshold of 50. These SNP calls were filtered out with the following criteria: MQ0Fraction > 0.001, QD < 4.3, within 6 bp of an indel, more than seven SNPs within a 100-bp region, Mapping Quality < 45, Homopolymer Run > 10, MQ0 > 9.5, Dels > 0.255. Moreover, only SNP calls covered by at least 5 non-reference reads across all libraries and 3 or more non-reference reads in at least one library were retained. SNPs that violated Hardy-Weinberg equilibrium with a binomial test P-value ($<1\times10^{-3}$) were removed as well. To eliminate mapping artefacts, SNPs in highly paralogous regions of the genome implicated by the ‘‘Self Chain’’ track on the UCSC Genome Browser [9] (normalized score R 90) were filtered out. Finally, a high-confidence set of 249,732 SNPs within H3K27ac peaks were obtained. We did not perform genotype calling since the G-SCI test does not require prior knowledge of genotype, instead it integrates over the likelihoods of all three genotypes for each individual, given the data [10]. Six samples were discarded when we discovered that they were incorrectly labelled as “European” using LASER **(Extended Fig. 10)** [11]. The remainder 64 samples were used for subsequent haQTL analysis.

**Identification of haQTL**

haQTLs were called in the 64 samples using G-SCI test [10]. This was done after all technical and biological covariates, importantly, disease etiology were regressed out. Top PCs which account for more than 5% variance were thus adjusted for. G-SCI test was performed on each of the 249,732 SNPs within peaks. For each SNP, an adjusted *P*-value was computed using a permutation test from 10,000 to 1 million permutations until a nonzero *P*-value was obtained. After 1 million permutations, if the adjusted *P*-value was still 0, it was set to $5x10^{-7}$. We then used the Benjamini and Hochberg multiple testing correction to calculate the FDR. At FDR threshold of 10%, 12,006 candidate haQTLs were identified. To detect possible artificial haQTLs due to different mapping rates to the reference genome between alleles, we simulated all possible 100 bp paired-end reads covering the haQTL and flanking SNPs and indels. The union of our SNP and indel (quality > 50 by GATK) calls and the 1000 Genome EUR SNPs and indels (1000 Genomes Project Consortium, 2012) were used. The fragment length of the simulated paired-end reads was set to be equal to 180 which is the median fragment size of all libraries. The simulated reads were then mapped to the reference genome using BWA. 4,231 haQTLs were discarded because their inferred allelic imbalances from the ChIP-seq data were smaller than five times the mapping bias estimated from the simulation. The remaining haQTLs were further filtered by an effect-size filter which calculated the Pearson correlation between peak height and the fraction of Q30 nonreference bases. haQTLs with $R^2 < 0.1$ were discarded. The final set of 1,680 haQTLs were in the remaining haQTLs after effect-size filter and only the most significant SNP (sentinel SNP) in each ChIP-seq peak was retained.

**PCM-1+ cardiomyocyte nuclei FAC sorting to determine cardiomyocyte cell purity**

50 mg of snap frozen human heart left ventricle tissue was cut and pounded into fine powder in liquid nitrogen. Tissues were rinsed in cold PBS (Takara, 1X protease inhibitor) and lyse in 10 ml cell lysis buffer (50mM HEPES-KOH, pH 7.5, 150mM NaCl, 1mM EDTA, 1% Triton X, 0.1% Sodium deoxycholate, 0.1% SDS, Takara 1X protease inhibitors) using a glass douncer with tight pestle ‘A’ (10-15 strokes). Crude nuclei were filtered through 70 µM and 40 µM cell strainers. Filtered nuclei were subjected to overnight incubation with cardiomyocyte-specific anti-PCM1 antibody (Sigma Aldrich, cat: HPA023374) at 1:1000 dilution. Nuclei were secondary stained with Goat anti-rabbit Alexa Fluor 488 secondary antibody (Thermofisher scientific, cat: R37116) at 1:500 dilution for 30 min. Nuclei were further stained with DAPI (Thermofisher scientific, cat: D9542) at 1:1000 dilution for 5 min before proceeding for FAC sorting on the BD biosciences Aria II sorter.

**RNA-seq library construction, sequencing and bioinformatics analysis**

Total RNA was extracted using the miRNeasy Kit (Qiagen) including DNAse treatment. RNA concentration and quality was determined using the NanoVue Plus™ spectrophotometer (GE Healthcare) and the Agilent 2100 RNA Nano Chip (Agilent). RNA sequencing libraries were prepared using the Illumina TruSeq stranded mRNA kit followed by the Nugen Ovation amplification kit. To avoid confounding by batch effects, libraries were randomly selected into pools of 32, and pools were sequenced on a Hiseq2500 to a depth of ~30 million 100 bp paired-end reads per biological sample. Fastq files were aligned against human reference (hg19/hGRC37) using the STAR aligner [12]. Duplicate reads were removed using MarkDuplicates from Picard tools, and per gene read counts for Ensembl (v75) gene annotations were computed. Expression levels in counts per million (CPM) were normalized and transformed using VOOM in the LIMMA R package [13].

**Expression-QTL (eQTL) analysis**

eQTL analysis with our LV samples was performed with linear regression analysis. The FPKM expression level was corrected for technical confounding factors including batch,

RIN, library size and fragment length, as well as biological factors such as etiology-related traits, sex, height and weight. PCA was performed again after regression to confirm that no confounding factors correlated strongly with the top 5 PCs. We considered cis-eQTL interactions with less than 100 kb separating the SNP and the TSS. We retained 180 eQTLs at FDR ≤ 20%. For distal-eQTL, we considered haQTL-gene pairs that were confined within the same TAD, and having minimal distance of 10 kb between haQTL and TSS. In order to refine the search for distal eQTL, only haQTL-gene pairs connected by HiChIP chromatin loops were considered.

**Human cardiac Hi-C**

*In situ* Hi-C was performed as previously described [14]. 70 mg of snap frozen human heart left ventricle tissue was cut and pounded into fine powder in liquid nitrogen. Tissues were washed twice in cold PBS, crosslinked in 1.5% formaldehyde for 10 minutes at room temperature after which glycine was added to stop the reaction. Cross-linked cells were lysed in cold lysis buffer (10 mM Tris-HCl pH8.0, 10 mM NaCl, 0.2% (v/v) Igepal CA630, mixed with protease inhibitors) on placed ice for 30 minutes. Subsequently, we applied 15 strokes using a glass douncer tight pestle ‘A’ to release nuclei. Pellets were washed twice with cold 1x NEBuffer 2, resuspended in 50 µl of 0.5% SDS and incubated at 62ºC for 10 minutes to solubilize the chromatin. After heating, 120 µl of water and 50 µl of 5% Triton X-100 (Sigma 93443) was added to quench the SDS. Chromatin was digested overnight by adding 25 µl of 10X NEBuffer 2 and 100U MboI (New England Biolabs, Cat#:R0147). Digestion efficiency was confirmed by isolating DNA from a 25 µl aliquot of the suspension and performing agarose gel electrophoresis. Digested ends were filled and labelled with biotin by adding 37.5 µl of 0.4 mM Biotin-14-dATP (Thermo Fisher Scientific 19524016), 1.5 µl of 10mM dCTP, 1.5 µl of 10 mM dGTP, 1.5 µl of 10 mM dTTP, 8 µl of 5U/ µl DNA polymerase I, large klenow fragment (New England Biolabs Cat#:M0210) and incubated at 37ºC for 90 minutes. Blunt end ligation was performed by adding 463 µl of water, 120 µl of 10X NEB T4 DNA ligase buffer, 200 µl of 5% Triton X-100, 12 µl of 10 mg/ml BSA and 5 µl of T4 DNA Ligase (New England Biolabs, Cat#:M0202), the solution was incubated at 16ºC for 4 hours. After ligation, crosslink was reversed overnight and DNA was purified using Phenol-chloroform method. Next, Hi-C libraries were treated with T4 DNA Polymerase (New England Biolabs, M0203) to remove biotin from unligated ends. To do this, 5 µg of Hi-C library was combined with 2.5 µl of 1 mM dATP, 2.5 µl of 1 mM dGTP, 10 µl of 10X NEBuffer 2.1, 5 µl of T4 DNA Polymerase (New England Biolabs, Cat#:M0203) and water was added to a final volume of 100 ul. The reactions were incubated at 20ºC for 4 hours and stopped by incubating at 75ºC for 20 minutes. Samples were sheared to a mean fragment length of 400 bp using Covaris M220. Pull-down of biotinylated DNA was performed using Dynabeads My One T1 Streptavidin beads (Thermo Fisher Scientific Cat#:65601), and library prep was performed following NEBNext® Ultra™ II DNA Library Prep Kit for Illumina®. Samples were quantified using KAPA and sequencing performed on a Hiseq 4000.

**Hi-C Analysis**

All sequencing QC are described in Supplementary Table 2. FASTQ sequences were mapped and filtered using HiCUP pipeline[15]. Hi-C reads (paired end, 2x151 bp) were aligned against the hg19 genome using Bowtie2. Only unique high-quality alignments were retained. Forward and reverse reads were mapped independently. Sequences which were mapped unambiguously to the genome were retained. Di-tags representative of experimental artefact were filtered out as these invalid di-tags might lead to inaccurate inference concerning genomic structure. These invalid di-tags were detected by positioning the putative di-tags on an in silico digested reference genome. Sequences which were mapped to the same restriction fragment were discarded as they did not describe the multi-dimensional contacts. Di-tags insert sizes which did not fit within the size-selection range were also discard. Lastly, PCR duplicates were removed. Cleaned-up aligned sequenced files in BAM format were processed by HiCPro [16] to generate raw interaction count at various bins of equal sizes (10 kb, 40 kb). Contact matrices with raw interaction counts were normalized using iterative correction of biased based on ICE algorithm[17]. TAD-calling was performed using Domaincaller[18], based on the ICE-normalized count matrix at 40 kb resolution. Fit-hi-C [19] was also used to identify significant chromatin interaction at 10 kb resolution. Insulation-scores were calculated as described in[20], using a window-size of 40 kb.

**Human heart HiChIP**

HiChIP was adapted from a previously published protocol [21] with some modifications. Briefly, 50 mg of snap frozen human heart left ventricle tissue was cut and pulverized in liquid nitrogen using mortar and pestle. Tissues were washed twice in cold PBS, and Hi-C portion performed following the Arima-HIC protocol described in the Arima Hi-C Kit (Product No A510008). For the ChIP portion, the nuclear pellet was sonicated to obtain chromatin fragments between 200 to 500 bp. Sheared chromatin was immunoprecipitated with 3 µg of H3K27ac antibody (Abcam, ab4729) with 30 µl of protein G beads (Invitrogen) overnight at 4ºC. Beads were washed and eluted in 200 µl of elution buffer (50 mM sodium bicarbonate, 1% SDS), de-crosslinked at 67ºC for 2 hours and DNA extracted using the Qiagen MinElute kit. Library preparation was performed using NEB ultra II library preparation kit, according to manufacturer's protocol. 10-12 PCR cycles were performed using indexed primers and libraries size between 300 to 500 bp were selected. The libraries underwent paired end sequencing at 2 x 151 bp read length on Illumina Hiseq 4K platform.

**HiChIP Analysis**

A similar analysis workflow from HiC above was applied to HiChIP. FASTQ sequences were mapped and filtered using HiCUP pipeline [15]. Filtered sequenced files in BAM format were analysed by HiCPro to generate raw interaction count. Contact matrices at 10 kb resolution were normalized using iterative correction of biased based on ICE algorithm[17]. Filtered sequenced files in BAM format were also converted into .hic files using pre function in juicer package[22]. High confidence chromatin loops were identified using the Juicer pipeline HiCCUPS tool at 5 kb and 10 kb resolution using the default parameters: hiccups –m 500 –r 5000,10000 –f 0.1,0.1, -p 4,2 –i 7,5, -d 20000,20000 <.hic> <output> [14]. Biological and technical reproducibility assessment between HiChIP samples were performed by comparing the total number of di-tags supporting each chromatin loop called from the merged replicates **(Extended Fig. 6a).** The pearson correlation between HiChIP replicates was computed from ICE-normalized count using the R cor() function.

#### Transcription Factor (TF) Motif Alteration Prediction

R package MotifBreakR [23] was used to identify transcription factor motifs that overlapped with the 1680 haQTLs using motif databases from Jaspar 2018 [24] using the default parameters: filterp=TRUE, pwmList=jaspar2018, threshold=1e-4, method="ic". The motifs of transcription factor binding was predicted to be significantly altered are listed in the Supplemental Table 10. Only TFs that were found to be expressed in the human RNA-seq samples were used for the analysis (FPKM>=1).

#### Intersection against published GWAS datasets

A total of 10 sets of GWAS datasets was downloaded [25-29]. LD ($R^2$>0.8) between QTLs and GWAS SNPs was calculated using genotypes from the European (EUR) 1000 Genomes. To show that haQTLs were enriched in GWAS datasets, permutation analyses were performed. Taking $n$ to be the number of overlaps between all SNPs in each GWAS dataset and the 1,680 haQTLs, we sampled 1,000 sets of $n$ random SNPs for each permutation analysis, matching for MAF and the distance to the nearest TSS. The expected number of overlaps between all 1,000 permutation sets and GWAS hits (above adjusted *P*-value < $1\ x\ 10^{-3}$) for each dataset was thus estimated. Fold enrichment was taken as the ratio of observed to expected.
